# Regularized logistic regression-based gene selection for functional analysis of heterogeneous scRNA-seq macrophage populations in muscle aging and regeneration

**DOI:** 10.64898/2026.10.25.761682

**Authors:** Tiago M. Costa, Marta B. Lopes, Joana Neves, Pedro Sousa-Victor, Susana Vinga

## Abstract

**Background:** Single-cell RNA sequencing enables the characterization of heterogeneous cellular responses across experimental conditions. However, feature-selection methods often prioritize genes associated with global condition differences while overlooking subgroup-specific transcriptional structure. This limitation is particularly relevant in complex tissues, where distinct cell states may contribute differently to disease, aging, or regeneration. Regularized logistic regression provides an interpretable framework for sparse gene selection, but standard penalties do not explicitly account for differences in gene-gene correlation patterns across annotated subgroups. Results: We introduce DisTwiner, a subgroup-aware extension of the twiner correlation-based regularization framework for binary classification problems with multiple annotated groups. DisTwiner computes pairwise angular scores between feature-correlation profiles across subgroup labels and incorporates the resulting feature-specific weights into an Elastic Net-regularized logistic regression model. We evaluated DisTwiner using simulated data and a single-cell RNA-seq dataset of macrophage populations isolated from young and aged regenerating murine skeletal muscle. In simulated data, alternative aggregation and transformation strategies showed comparable classification performance, with minimum-based exponential weighting producing the greatest dispersion of feature penalties. In the macrophage dataset, DisTwiner achieved classification performance comparable to standard Elastic Net logistic regression and higher than Random Forest under the evaluated setting. Although predictive accuracy was similar to Elastic Net, DisTwiner selected genes with more heterogeneous fold-change and correlation patterns across macrophage subpopulations. Downstream enrichment analyses showed that DisTwiner-selected features captured biologically relevant aging-associated pathways, including interferon-response signatures in specific macrophage populations, and could serve as an additional filtering layer after differential expression analysis. Conclusions: DisTwiner extends correlation-based regularization to binary classification settings with multiple annotated subgroups, enabling feature selection that accounts for subgroup-specific correlation structure. Applied to single-cell macrophage data, the method preserved competitive classification performance while prioritizing genes with cell-population-dependent expression patterns, supporting its use for interpretable subgroup-aware feature selection in high-dimensional omics data.

## 1 Introduction

Regeneration is a biological process essential for the maintenance of multicellular organisms, in which damaged or disturbed tissues are restored by the coordinated action of the remaining viable cells in the organism [1]. An essential set of cellular functions preserved among different organisms in tissue regeneration involves the supportive role of multiple immune cell types infiltrating the damaged area, facilitating the clearance of necrotic debris caused by the injury, and managing the activities of other cells within the niche, a dynamic that is compromised in aging [2, 3, 4]. Aging is a multifactorial process associated with compromised functional integrity of tissues and organs, with decreasing maintenance and regeneration potential that affects cell types differently [5]. This phenomenon, while extensively acknowledged by human civilization at the level of its culture, history, and more recently scientific advances, is still lacking a clear understanding of the biomolecular mechanisms and communication axes involved in aging [6].

The adult skeletal muscle is a widely acknowledged model of regeneration in mammals, owing in great part to a specialized stem cell compartment, the satellite cells, that interacts with immune-derived signalling and has the potential to differentiate into myocytes that can then fuse to give rise to new myofibers, fully restoring the skeletal muscle in homeostatic conditions [7]. Immune regulation is essential for satellite cell function during regeneration [8], therefore exploring the extent of the functional heterogeneity of the immune cells in the skeletal muscle and how it varies with aging is a crucial step towards understanding the age-related loss of regeneration potential.

The emergence and rising popularity of single-cell RNA sequencing (scRNA-seq) technology has led to considerable improvements in modern biology research, enabling the profiling of transcriptomic alterations in biological samples at single-cell resolution [9]. Such breakthrough has enabled the development of new methods for the study of cellular dynamics such as trajectory inference, RNA velocity, and computational intercellular communication modelling [10, 11, 12]. In particular, intercellular communication analysis is an important tool to build insight into the complexity of regulatory networks within tissue niches across a plethora of cellular dynamics, such as cancer development, aging, and regeneration [13, 14].

Given the nature of scRNA-seq data, most datasets consist of thousands to millions of observations (the isolated cells), each with a number of features (the counts for each gene) of the order of tens of thousands, therefore, most scRNA-seq data are characterized by high dimensionality [15], which hinders classification of data groups based on traditional distance metrics and statistical learning models. With these limitations in mind, there is considerable interest in effectively identifying the features that contribute the most to the variability of the data. In the case of scRNA-seq data, this allows us to focus on the genes that are most likely to be relevant to a given biological question.

Machine learning (ML) approaches, particularly supervised models, provide a natural framework for combining feature selection with predictive modeling, enabling the identification of informative genes while simultaneously optimizing classification performance. In order to optimize classification using supervised ML models, we can use feature selection strategies that alongside classification prioritize the most explanatory features, thus improving classification by minimizing noise in the data [16, 17]. Popular feature selection methods for scRNA-seq include differential gene expression tests [16]. In a binary classification problem, these tests typically assign most observations to one of the two experimental classes. However, in many real datasets, there may be additional latent labels that further subdivide observations within each class. In particular, for scRNA-seq, feature selection methods that can isolate both within-class heterogeneity and coherent patterns across conditions are specially promising, as class-specific labels may represent different cell types, each exhibiting its own pattern of biomarker changes during aging, and detecting such patterns allows the refining of our understanding of complex biological dynamics.

Many of the feature selection methods currently available in the literature, including ML models of logistic regression with regularization terms, such as LASSO and Elastic Net [18], and differential expression tests such as MAST [19], do not explicitly account for labels beyond the primary experimental condition. As a result, they cannot automatically distinguish between biomarkers that show consistent variation across cell types and those with unique, cell-type-specific patterns, often requiring manual curation of the selected features. To address this limitation, it would be beneficial to develop a feature selection method that can handle an arbitrary number of labels in the data, thereby automating and streamlining the identification of biomarkers that vary either coherently or distinctly across cell types.

To capture consistent gene expression patterns across classes, Lopes et al. (2019) proposed `twiner`, a feature selection framework for classification that combines logistic regression with correlation-based regularization to prioritize genes exhibiting similar correlation patterns across experimental conditions [20]. The `twiner` model computes for each data label, such as disease or cell type, the covariance matrix of all features in the dataset, in which each column (or row) of the matrix represents the covariance vector a feature with the remaining ones. Model fitting is then performed using the experimental conditions, such as healthy versus diseased tissue, and each feature is penalized based on a dissimilarity measure given by the angle formed between the two respective covariate vectors of the same feature, taken in the two different labels or groups. The `twiner` regularizer can be used for the selection of features that are common to both labels used, in the case we penalize higher dissimilarity, but it is also possible to instead identify label-specific features by penalizing lower dissimilarity score [21]. However, given the nature of the dissimilarity metric, `twiner` is currently limited to only two labels in the entire dataset, whereas in the context of single-cell analysis clustering strategies are usually capable of identifying more than two cell types in the data.

In this study, we benchmark different classification and feature selection ML models in the context of scRNA-seq data and propose DisTwiner, an extension of the `twiner` regularizer for correlation-based feature selection that can accommodate an arbitrary number of sample labels across the conditions of the experimental design of a binary classification problem. We consider a scRNA-seq dataset of macrophages populating the murine skeletal muscle during regeneration in young and aged mice, where each macrophage is labeled as one of six possible functional profiles according to macrophage marker expression identity (Sousa et al., 2024) [2]. Our goal is to use a selection of ML models to identify biomarkers in the dataset associated with alterations of specific macrophage populations from young to aged mice, and compare the DisTwiner performance to that of gold standard strategies of feature selection and classification. The identified features can then be used as input for functional analysis specific to each macrophage population to improve our understanding of specific biological processes in aging and regeneration.

## 2 Methods

### 2.1 Transcriptomic data

#### 2.1.1 Synthetic data

To benchmark and optimize the DisTwiner pipeline, we generated a synthetic dataset organized into three groups *G* = {*G*_1_, *G*_2_, *G*_3_} observed under two experimental conditions *c ∈* {*i, ii*}. For each combination (*G*_*k*_, *c*), samples were drawn from a multivariate normal distribution

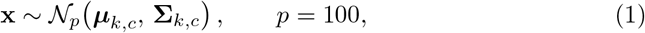

where *p* denotes the number of features, ***µ***_*k,c*_ *∈* ℝ^*p*^ is the mean vector, and **Σ**_*k,c*_ *∈* ℝ^*p×p*^ is the covariance matrix associated with group *G*_*k*_ under condition *c*.

The mean vectors were defined as follows. For group *G*_1_, entries of the mean vector were independently sampled from a uniform distribution, for each condition:

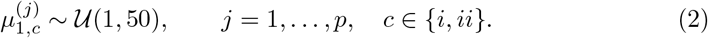

To induce correlation between groups in the simulated data, the mean vector of group *G*_2_ was obtained through a linear transformation of the mean vector of *G*_1_:

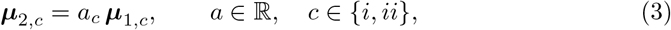

where 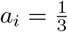 and 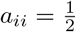.

For group *G*_3_, the mean vector was independently drawn from a standard normal distribution, ensuring a statistical profile distinct from *G*_1_ and *G*_2_:

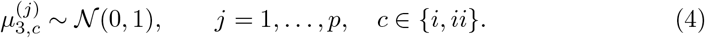

Covariance matrices **Σ**_*k,c*_ were constructed as positive definite matrices whose entries were sampled from a uniform distribution. Specifically, an auxiliary matrix **A** *∈* ℝ^*p×p*^ was generated with entries *A*_*ij*_ *∼ U* (0, 1), and the covariance matrix was obtained as

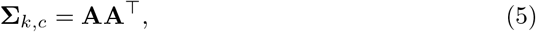

which guarantees positive definiteness by construction.

The final synthetic dataset consisted of *n*_1_ = 3500 samples for group *G*_1_ and *n*_2_ = *n*_3_ = 3000 samples for groups *G*_2_ and *G*_3_, respectively, introducing a mild class imbalance to better mimic realistic experimental scenarios.

#### 2.1.2 Experimental data

For experimental validation of the methods, we used the Muscle Immune Aging scRNA-Seq dataset (Sousa et al., 2024) [2]. The Muscle Immune Aging dataset was sequenced from a library of immune cells isolated from both young and aged murine skeletal muscle in homeostatic conditions and at different time points of regeneration, and enriched prior to library preparation using fluorescence-activated cell sorting (FACS) to separate CD45^+^ (immune) cells from the remaining cells in the skeletal muscle. In this study, we took into consideration the heterogeneity of one of the most abundant immune cell types during regeneration, the macrophages, as such we used the available recluster of the macrophage samples and the respective functional annotation provided in the original study [2]. The final dataset that we used for this study consisted of the log1p-normalized data, where each entry in the gene expression matrix *g*_*ij*_ is defined as:

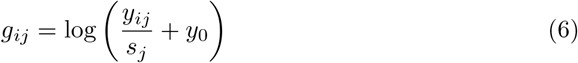

where *y*_*ij*_ denotes the associated raw count value of gene *i* in cell *j, s*_*j*_ is the library size factor for cell *j*, and *y*_0_ is the arbitrarily small pseudocount value, provided by the `Seurat R` library. The resulting gene expression matrix consists of 23,501 cells across 16,864 genes/features. Macrophages, both from young and aged samples, were divided into one of six categories: Infiltrating macrophages (infMacs), regulatory-associated macrophages 1 and 2 (regMacs 1 and 2, respectively), intermediate macrophages (intMacs), repair-associated macrophages (repMacs), and late repair-associated macrophages (lateMacs).

### 2.2 Sparse Classification

In this study we focused on sparsity-based solutions for biomarker identification in scRNA-Seq data with likelihood-based classification. We addressed a binary classification problem aimed at identifying young versus aged cells (condition) within the dataset, wherein each cell also possesses a secondary label assigning it to one of six clusters (group). We employ DisTwiner as a feature selection model for the identification of genes with differing transcriptomic alteration patterns across groups in the data. As a baseline for comparison with DisTwiner, we used logistic regression models. Logistic regression fits a sigmoid curve to estimate the probability of class membership and was combined with an Elastic Net regularization term in the cost function to promote feature selection, yielding sparse solutions that discards non-explanatory features [22].

#### 2.2.1 Regularized logistic regression

*Binary logistic regression* describes the relationship between one or more features and a binary outcome, which is given by the logistic function:

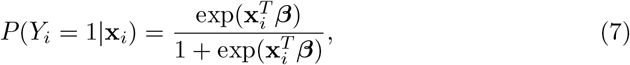

where *P* (*Y*_*i*_ = 1 |**x**_*i*_) is the probability of ‘success’ for observation *i*, or the probability of belonging to a given class.

The parameters of the logistic model are estimated by maximizing the log-likelihood function, given by

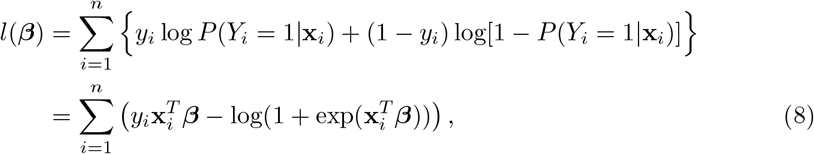

where the binary variable *y*_*i*_ indicates success (*y*_*i*_ = 1) or unsuccess (*y*_*i*_ = 0) for observation *i*. By the introduction of a regularization term, the log-likelihood function becomes

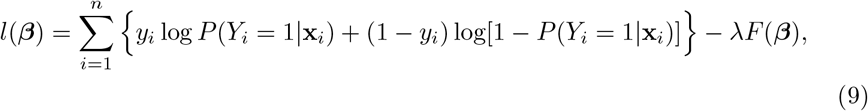

with

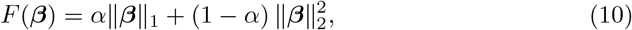

where *α* ∈ [0, 1] determines the balance between the LASSO (*α* = 1) and Ridge (*α* = 0) penalties and the tuning parameter *λ* controls the overall strength of regularization.

#### 2.2.2 Generalized DisTwiner formulation for an arbitrary number of groups

We initially define DisTwiner as an expansion of the twiner model that was recently introduced as a structured regularizer that leverages pairwise correlations between features belonging to two separate groups, *A* and *B* [20, 21], demonstrating the potential of regularization methods that explicitly incorporate the *correlation structure* of the data.

Let the training data be partitioned into *K* ≥ 2 latent groups (e.g., cell types or sample sub-labels) G = {*G*_1_, *G*_2_, …, *G*_*K*_}, each observed across the two experimental conditions used for the binary classification task. For every group *G*_*k*_ ∈ *G* we compute the feature correlation matrix

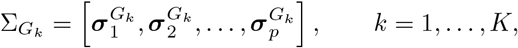

where 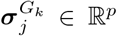 is the correlation vector of feature *j* with all remaining features within group *G*_*k*_.

For dissimilarity metrics, we propose two different DisTwiner modes, one that identifies label-specific feature correlation profiles with larger discrepancies between labels (DisTwiner-specific, Eq. 11), and one that can isolate label-shared feature correlation profiles with minimal differences (DisTwiner-common, Eq. 12).

Extending the `twiner` rationale, we define the pairwise dissimilarity of feature *j* between any two groups *G*_*a*_, *G*_*b*_ *∈ G* in DisTwiner-specific mode as a normalized angular distance:

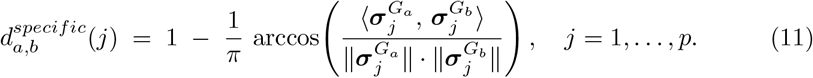

And define the dissimilarity of feature *j* between any two groups *G*_*a*_, *G*_*b*_ *∈ G* in DisTwiner-common mode as:

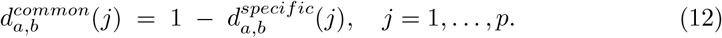

Let *P* (*G*) = {(*G*_*a*_, *G*_*b*_) |*a < b, G*_*a*_, *G*_*b*_ *∈ G*} denote the set of all 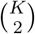 unordered group pairs. For each feature *j* we build the distribution of pairwise scores

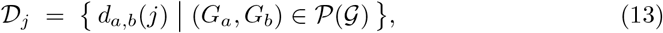

and summarize it through a summary statistics (mean, maximum, or minimum) to obtain the per-feature weight:

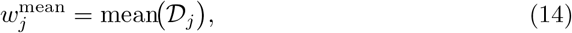

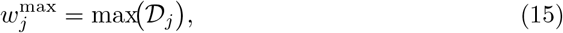

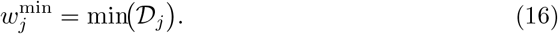

For benchmarking distributions we used a distribution of angles directly computed from Eq. 11 and 12, a distribution of an exponential transformation of the values (Eq. 17), and a distribution of a sigmoid transformation of the values (Eq. 18), resorting to non-linear transformations of the pairwise scores that can further tune the separation between features. Keeping the minimum-based aggregation (which yielded the best trade-off between accuracy and weight dispersion), we define:

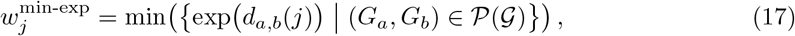

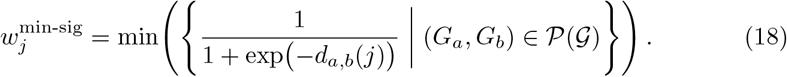

Finally, the per-feature weights are collected into **w** = (*w*_1_, …, *w*_*p*_) and plugged into the same Hadamard-weighted elastic-net penalty introduced for `twiner`:

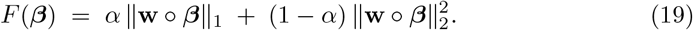

For *K* = 2, the model reduces (up to the 1 − (·)*/π* normalization) to the original `twiner` dissimilarity so DisTwiner is a strict generalization of `twiner` to *K* ≥2 latent groups.

We performed classifier performance evaluations by using samples from all six groups available in the dataset and separating the data across different folds into train subsets composed of randomly selected data points amounting to 80% of the total data, which was also used to compute the correlation matrices of each label, and test subsets built with the remaining 20%.

We used the synthetic data to assess the classification performance of each of the models differing by the central tendency metric with standard classifier evaluation metrics (accuracy, precision, recall, ROC AUC, and F1), as well as measuring the standard deviation of the final coefficients of each feature to evaluate which models produce higher degree of separation in the feature penalization. We then consider the model with best classification performance across all metrics, which turned out to be the minimum-based one, and once again benchmarked classification performance, this time varying the type of distribution.

We then evaluated the classification performance in real transcriptomic data against a selection of established classification models (Elastic Net-regularized logistic regression and Random Forest) using standard metrics for machine learning classification evaluation (accuracy, precision, recall, ROC AUC, and F1). We optimized the Elastic Net-regularized logistic regression classifier using 10-fold cross-validation using the `glmnet` library and modeled the Random Forest classifier with 500 decision trees using the `randomForeste` library, both available in R.

The overall pipeline is described in Figure 1.

**Fig. 1.**
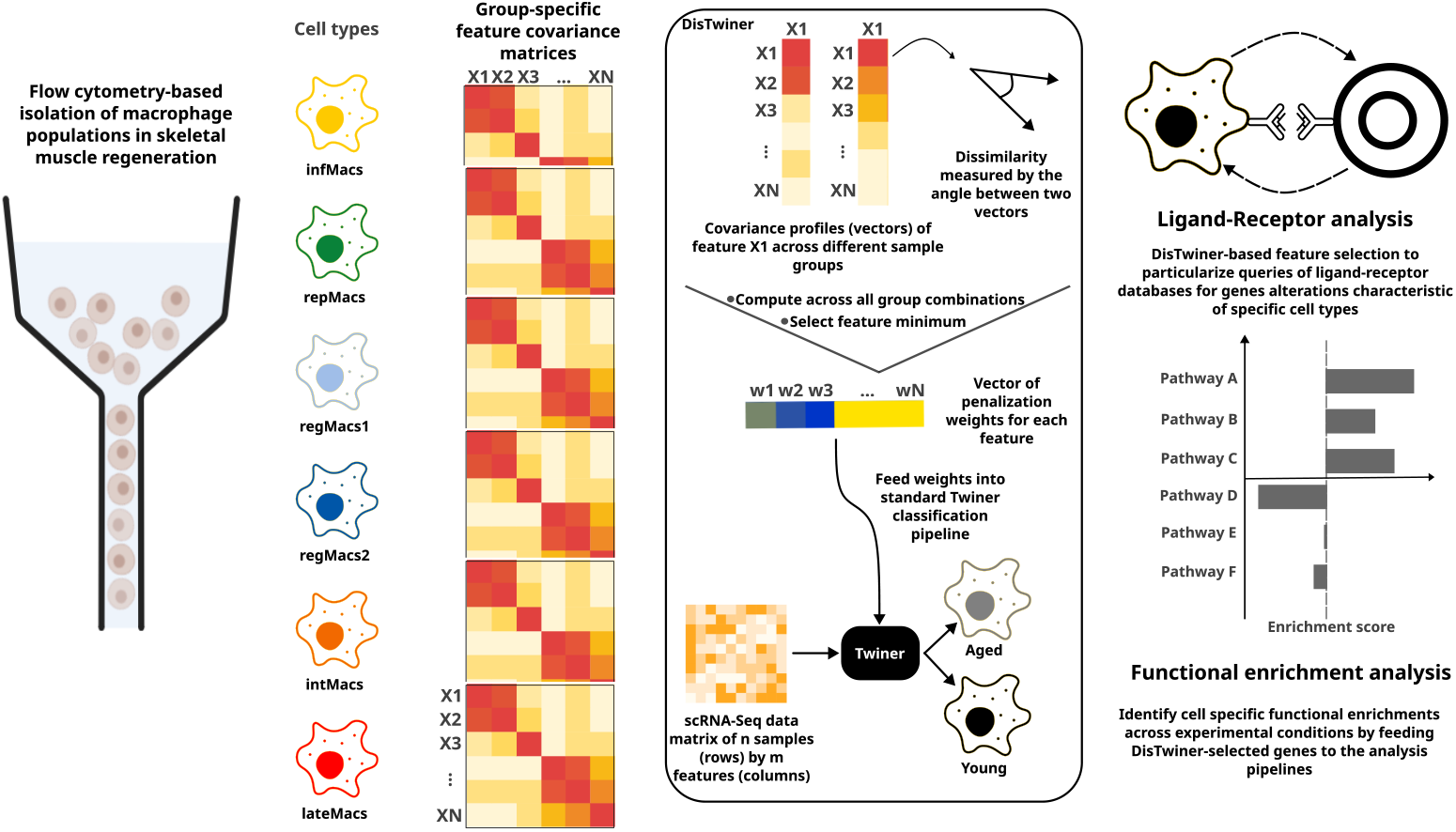
Summary of DisTwiner workflow. The data is split along experimental conditions and, orthogonally, along latent groups that span both conditions. For each group pair, a feature dissimilarity matrix is generated, from which we compute the penalization coefficient for each feature used in the regularization term of model fitting.

### 2.3 Functional validation of DisTwiner-based feature selection

Following model training for classification of young and aged samples, we compared the overlap of features selected by the DisTwiner model with features selected by standard Elastic Net-regularized logistic regression trained with the Muscle Immune Aging data, and the features obtained from the MAST differential expression test (we selected the top 2000 features on the basis of the p-value, with Benjamini-Hochberg correction) found in the implementation of the `Seurat` R package. We then performed Gene Set Enrichment Analysis (GSEA) with the `fgsea` R package for all sets of selected features using the log fold-change across the two conditions as the score of each feature, and compared the overlap of Hallmark and Reactome pathways found for each method. We selected pathways with absolute Normalized Enrichment Score ≥ 1 and defined statistically significant enrichments with adjusted p-value ≤ 0.05.

## 3 Results

### 3.1 DisTwiner correctly classifies simulated data composed of multiple groups with different distributions

In order to optimize the distance metric of the `twiner` model, we computed angles between the feature correlation vectors of a feature between two groups for all available group combinations and then selected a central tendency metric to represent the distribution of the given feature. We computed the distribution of penalization factors obtained based on the mean, maximum, and minimum of the distributions of each individual feature, and observed varying degrees of dispersion in the resulting distributions (Figure 2.A-C).

**Fig. 2.**
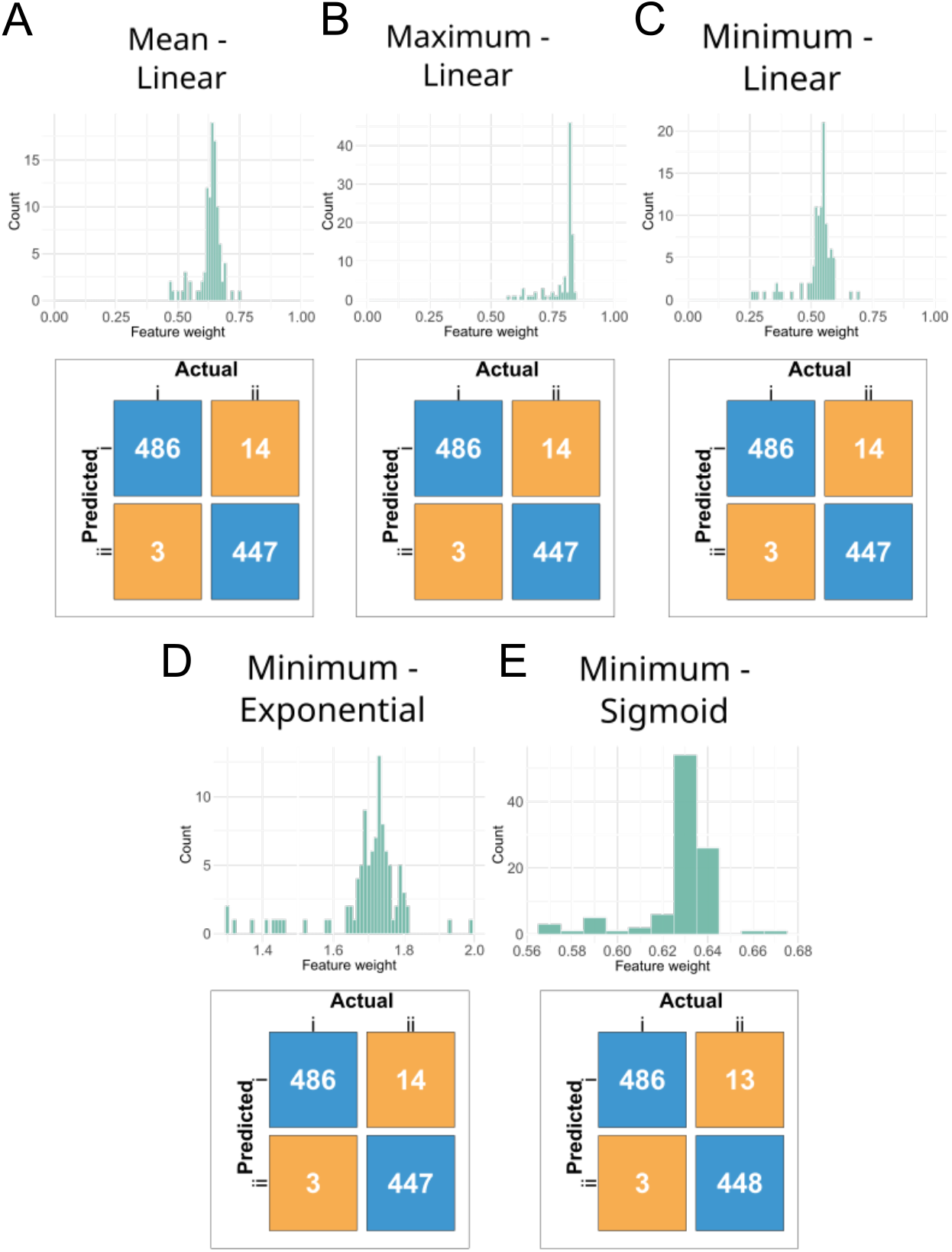
DisTwiner benchmarking and optimization using simulated data. **(A)** Histogram of feature weights obtained from the mean of the coefficients computed across all possible pairs of groups in the data for every single feature and the confusion matrix for the respective classification benchmark. **(B)** Histogram of feature weights obtained from the maximum of the coefficients computed across all possible pairs of groups in the data for every single feature and the confusion matrix for the respective classification benchmark. **(C)** Histogram of feature weights obtained from the minimum of the coefficients computed across all possible pairs of groups in the data for every single feature and the confusion matrix for the respective classification benchmark. **(D)** Histogram of feature weights obtained from the exponential transformation of the minimum of the coefficients computed across all possible pairs of groups in the data for every single feature and the confusion matrix for the respective classification benchmark. **(E)** Histogram of feature weights obtained from the sigmoid transformation of the minimum of the coefficients computed across all possible pairs of groups in the data for every single feature and the confusion matrix for the respective classification benchmark.

We then evaluated each classification model using standard metrics and the standard deviation of the distribution of coefficients as an estimate of dispersion (Table 1).

**Table 1.** Comparison metrics between different strategies of weight computation of the DisTwiner model.

| Model | Accuracy | Precision | Recall | F1 | ROC<br>AUC | Weights<br>std. dev. |
| --- | --- | --- | --- | --- | --- | --- |
| Mean Linear | 0.970±0.005 | 0.973±0.006 | 0.965±0.009 | 0.969±0.005 | 0.997±0.001 | 0.049±0.001 |
| Max Linear | 0.970±0.004 | 0.973±0.006 | 0.964±0.008 | 0.969±0.005 | 0.997±0.001 | 0.067±0.001 |
| Min Linear | 0.970±0.005 | 0.973±0.006 | 0.965±0.009 | 0.969±0.005 | 0.997±0.001 | 0.075±0.001 |
| Min Exp | 0.970±0.005 | 0.973±0.006 | 0.965±0.009 | 0.969±0.005 | 0.997±0.001 | **0.118±0.001** |
| Min Sig | 0.970±0.005 | 0.973±0.006 | 0.964±0.008 | 0.969±0.005 | 0.997±0.001 | 0.018±0.001 |

**Table 2.** Classification performance metrics of DisTwiner classifier and selected standard classification models on the Muscle Immune Aging scRNA-Seq dataset.

| Model | Accuracy | Precision | Recall | F1 | ROC AUC |
| --- | --- | --- | --- | --- | --- |
| DisTwiner | 0.932±0.005 | 0.934±0.005 | 0.929±0.006 | 0.932±0.005 | 0.982±0.002 |
| Elastic Net-regularized Logistic Regression | **0.933**±0.005 | 0.935±0.004 | **0.931**±0.007 | **0.933**±0.005 | 0.982±0.002 |
| Random Forest | 0.907±0.002 | **0.937**±0.007 | 0.872±0.009 | 0.903±0.003 | 0.973±0.001 |

We observed good classification performance amongst all tested models as based on the standard evaluation metrics, and observed variations in the dispersion of the feature weights after applying the central tendency metric to each distribution across metrics. We consider this dispersion to be relevant in predicting the classification performance of these models in real biological data, as the aim of DisTwiner will be to identify features with strong correlation/dissociation between groups in the classification of the hypothetical experimental conditions *i* and *ii*, and as such a higher variability in the weights used for feature selection may improve the separation and removal of features redundant for classification. Based on this assumption, we selected the model that considers the minimum of the distribution for subsequent evaluations. We then sought to explore variations in distance computation by benchmarking the linear score (Eq. 16) against nonlinear transformations of the values (Figure 2.D-E) by computing exponential and sigmoid functions of the angle between each vector (Eq. 17 and 18). We compared the performance of the three metrics based on the evaluation parameters described above (Figure 2.D-E and Table 1). We observed a similar classification accuracy between the total of five models (Table 1); however, the sigmoid transformation model showed a decrease in the dispersion of the feature weights while the exponential transformation model exhibits an increase of approximately 60% in the standard deviation of feature weight (Table 1); therefore, in the course of this study we selected the model described by Eq. 17 for classification benchmarking with real data and interpretation of the biological significance of the feature selection output.

### 3.2 DisTwiner classifies scRNA-Seq data between two experimental conditions across multiple transcriptionally distinct cell clusters

We then applied the DisTwiner classification model to biological data. We used the Muscle Immune Aging scRNA-Seq dataset [2], which identifies a high number of heterogeneous macrophage populations (Fig. 3.A) that we used to evaluate the performance of the model in classifying whether a cell is young or old across multiple cell clusters.

**Fig. 3.**
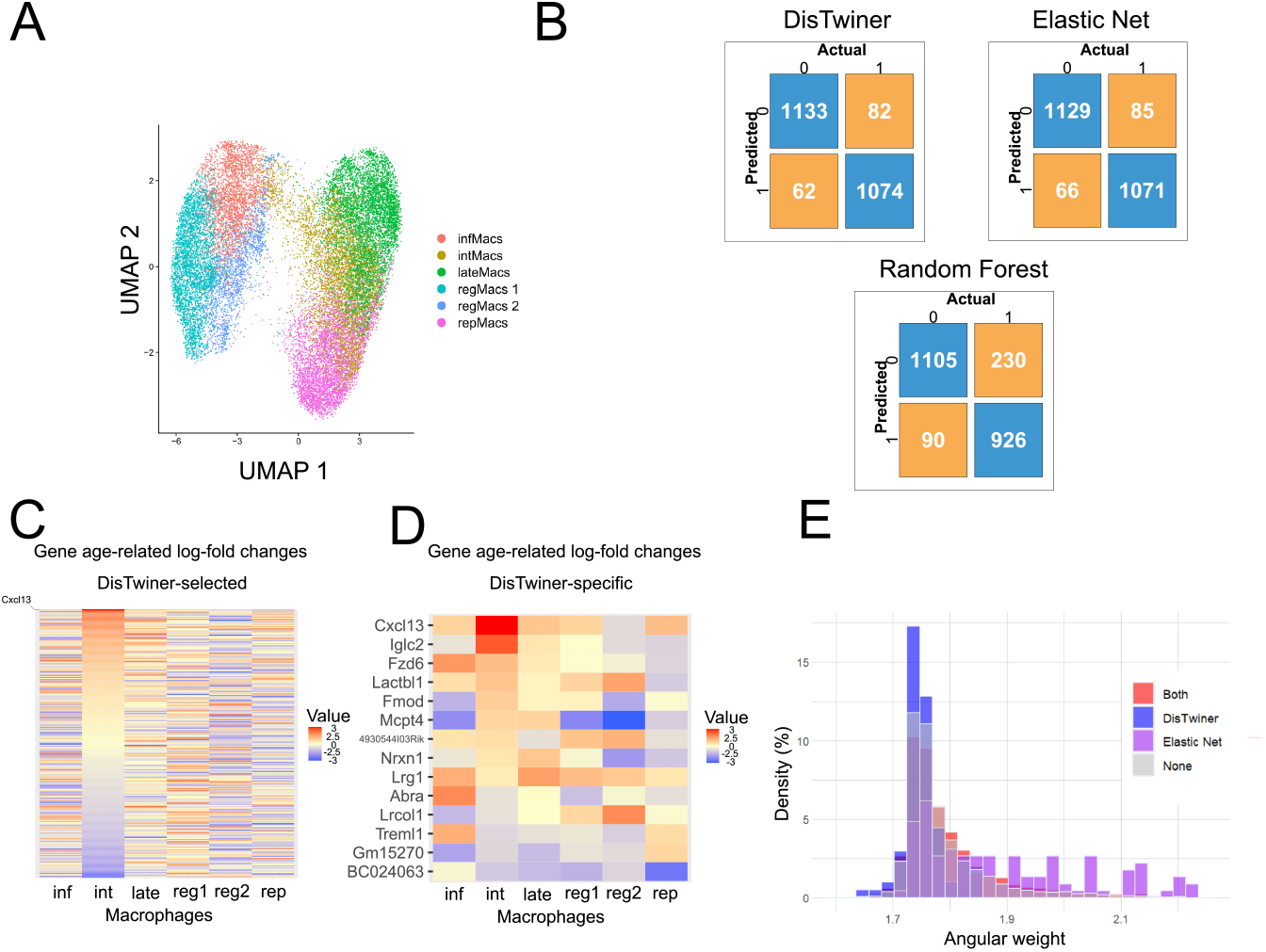
DisTwiner benchmarking using real data exhibits satisfactory performance in the classification task for scRNA-Seq data. **(A)** UMAP plot of the macrophage recluster of the Muscle Immune Aging scRNA-Seq dataset at 0.6 clustering resolution used to benchmark the DisTwiner performance against standard classification models. **(B)** Confusion matrix of the young/old classification performance for the DisTwiner, Elastic Net-regularized logistic regression, and Random Forest models. **(C)** Heatmap of fold-change values of each feature selected by DisTwiner for each cell cluster. **(D)** Heatmap of fold-change values of each feature specifically by DisTwiner for each cell cluster. **(E)** Histogram of feature weights selected for each feature separating them by selection status relative to DisTwiner and Elastic Net.

We used the data including all available macrophage populations during regeneration (Fig. 3.A) and applied the DisTwiner method to compute distance scores for each feature. We observed that DisTwiner preserves classification robustness even with an increase of possible groups, with a classification accuracy of approximately 93% (Table 3). We then compared the accuracy of DisTwiner on the macrophage data with selected state-of-the-art classification models, namely Elastic Net-regularized logistic regression and Random Forest. We observed that logistic regression has high classification accuracy on the dataset that is on par with DisTwiner accuracy, whereas Random Forest displays a slight loss in accuracy (Figure 3.B; Table 3). These results suggest that logistic regression is an appropriate model for binary classification of young and aged cells in scRNA-Seq data, and that DisTwiner is capable of particularizing the feature selection to genes that may represent characteristic changes of a given cell type without compromising classification robustness.

**Table 3.**
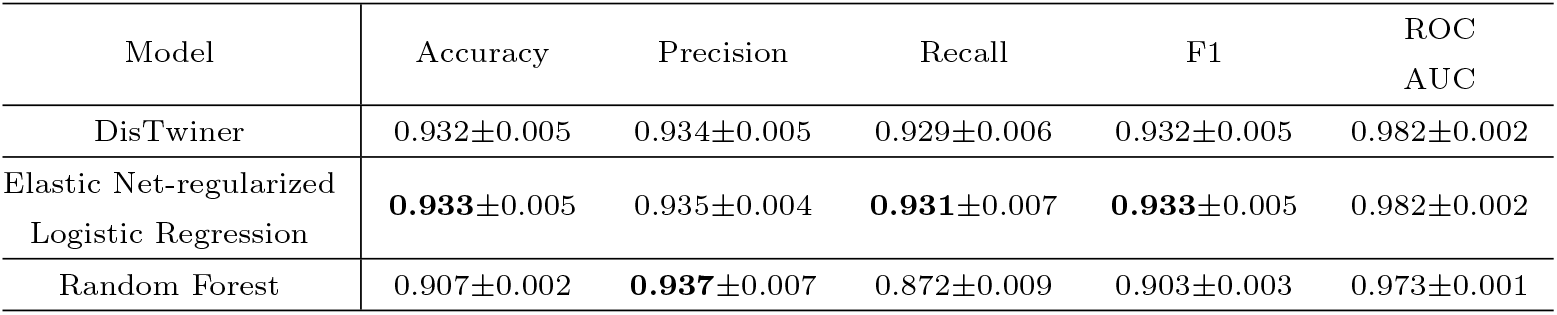
Classification performance metrics of DisTwiner classifier and selected standard classification models on the Muscle Immune Aging scRNA-Seq dataset.

We looked into the genes selected by the DisTwiner classification (Supplementary Data 1) and observed that the expression fold-change values across all macrophage groups for the majority of features varies substantially (Figure 3.C). Many features only exhibit alterations in a subset of macrophage clusters or entirely opposite fold-change patterns from one cluster to another (Figure 3.C). In particular, upon inspecting the subset of features selected exclusively by DisTwiner, we observed a distinct upregulation of the cytokine Cxcl13 on intMacs accompanied by downregulation of this gene in regMacs 2, a type of macrophage shown in the context of organ transplantation and cancer models to have immune-suppressive activity regarding the lymphocyte compartment [23, 24] (Figure 3.D). Cxcl13 is a cytokine responsible for promoting lymphocyte recruitment, therefore a possible interpretation is that aged intMacs upregulate Cxcl13 expression resulting in the observed age-related increases of T-cell recruitment [2]. We additionally compared the features selected by DisTwiner and Elastic Net on the basis of the penalization weights computed in the DisTwiner pipeline, and observed that Elastic Net tends to select features with higher coefficient whereas DisTwiner favors features with lower weights (Figure 3.E). These results suggest that DisTwiner feature selection is selecting genes with low penalization weight, as intended, with distinct correlation profiles from each other, supporting the hypothesis that DisTwiner is able to select features with varying correlation profiles across the samples.

### 3.3 DisTwiner feature selection identifies a reduced set of genes in comparison to differential gene expression tests

We sought to assess the biological significance of the genes selected by DisTwiner classification of young and aged cells by utilizing the features as input of functional analysis pipelines. We first plotted the overlap of genes selected by the DisTwiner classifier, Elastic Net, and the MAST differential expression test [19] for the repMacs cluster (Fig 4.A). We observed features uniquely selected by each of the methods, as well as a subset of common features selected by both methods (Fig 4.A), however MAST identified a considerably higher number of unique genes. We considered the implications of this reduced number of identified features in the DisTwiner set for downstream functional analysis by running Gene Set Enrichment Analysis (GSEA) with the genes selected by each method. Out of the total 50 pathways from the Hallmark collection, we observed that MAST-selected features yield 37 pathways above enrichment cut-off values for repMacs, and both DisTwiner and Elastic Net-selected features yield 7 additional pathways on top of the ones already identified using MAST (Fig 4.B). However, we observed that MAST is capable of identifying a higher number of pathways with statistically significant enrichment in the Hallmark collection, with DisTwiner features resulting in a reduced pool of statistically significant pathway enrichments, and no statistical significance for any gene set using Elastic Net-selected features (Fig 4.B).

**Fig. 4.**
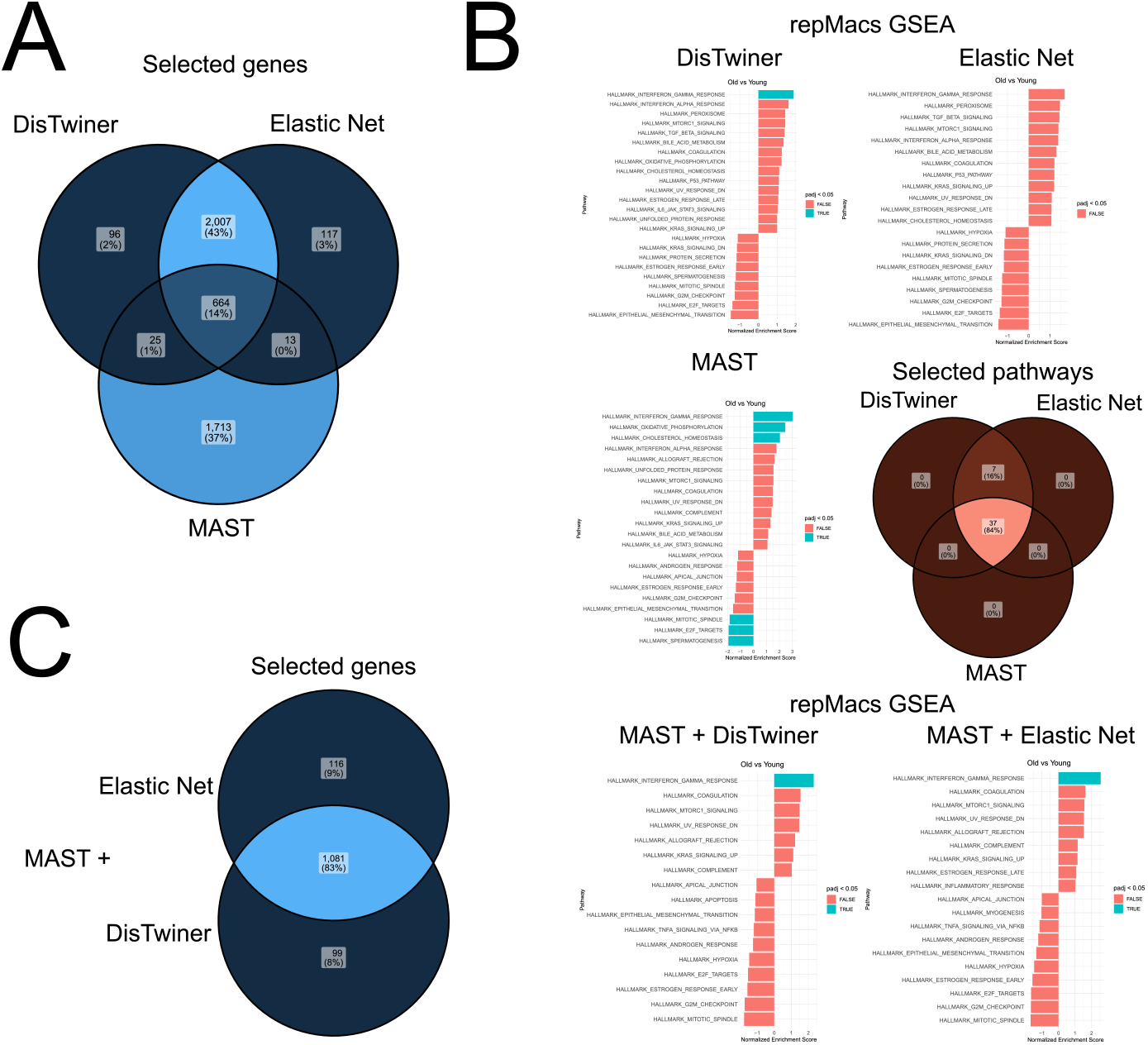
DisTwiner feature selection highlights unique genes and functional pathways in scRNA-Seq data. **(A)** Venn diagram of the overlap of features selected by DisTwiner, Elastic Net, and the MAST differential gene expression test for repMacs. **(B)** Plot of Hallmark pathways GSEA scores for the repMacs group using log-fold change from the selected features by each model comparing cells from aged and young samples as feature ranks alongside the Venn diagram showing the overlap of pathways selected by each method. **(C)** Venn diagram of the feature selection overlap and respective Hallmark pathways GSEA scores for the repeated analysis using features initially prefiltered by MAST.

We then considered that DisTwiner feature selection could be capable of improving the downstream functional analysis of differential gene expression tests such as MAST, by fitting the model based on data variability of a pre-selected list of differentially expressed genes (DEGs) by a highly popular method of feature selection in transcriptomic analysis, increasing the robustness of the features selected at the end of the pipeline. We performed GSEA with either DEGs directly obtained from MAST or only DEGs also selected by DisTwiner/Elastic Net, using the respective log-fold change values of the repMacs cluster as GSEA ranks Fig (4.C). We observed that the MAST DEGs allow the detection of several statistically significant enrichments using GSEA, such as positive enrichments of IFN-*α*/*γ* responses, as well of oxidative phosphorylation (Fig 4.C). When we provide the MAST DEGs + DisTwiner features to the GSEA pipeline instead, we still observe preserved functional enrichments, such as that of IFN-*γ*, but we also observe a narrowed down list of enriched pathways conserved from the original list on MAST DEGs such as the positive and negative enrichments of Kras signaling and androgen response, respectively (Fig 4.C). However, GSEA analysis of MAST DEGs + Elastic Net features yields a similar result to that of MAST DEGs + DisTwiner. These results suggest that while DisTwiner classification can provide an additional layer of robustness to the feature selection workflow by providing an additional criteria to satisfy, the difference in selected features does not consistently translate into an improvement in the identification of robust cell type-specific functional enrichments.

We then inspected the DisTwiner performance at identifying unique pathway enrichments across different cell types by comparing the GSEA results for each macrophage cluster using the features selected by DisTwiner (Figure 5). We observed that pathways identified with DisTwiner-selected features differ based on the cluster of choice, with infMacs and regMacs 2 outputting no significantly enriched pathway while regMacs 1 highlights a negative enrichment of the inflammatory response in aging (Figure 5). The results also show similarities between the enrichments of repMacs and lateMacs characterized by a positive enrichment of the IFN-*γ* response, while the intMacs are additionally characterized by a positive enrichment of the IFN-*α* response (Figure 5). Additionally, our results appear to suggest that beyond the enrichment profile of each individual cluster, the DisTwiner-selected features also identified pathway enrichments that separate clusters based on more general subsets such as Ly6C^high^ macrophages (infMacs, regMacs 1, and regMacs 2) [2] exhibiting an agerelated tendency for the negative enrichment of inflammatory responses while Ly6C^low^ macrophages (intMacs, repMacs, and lateMacs) [2] show a tendency for age-related increases in IFN response activity (Figure 5).

**Fig. 5.**
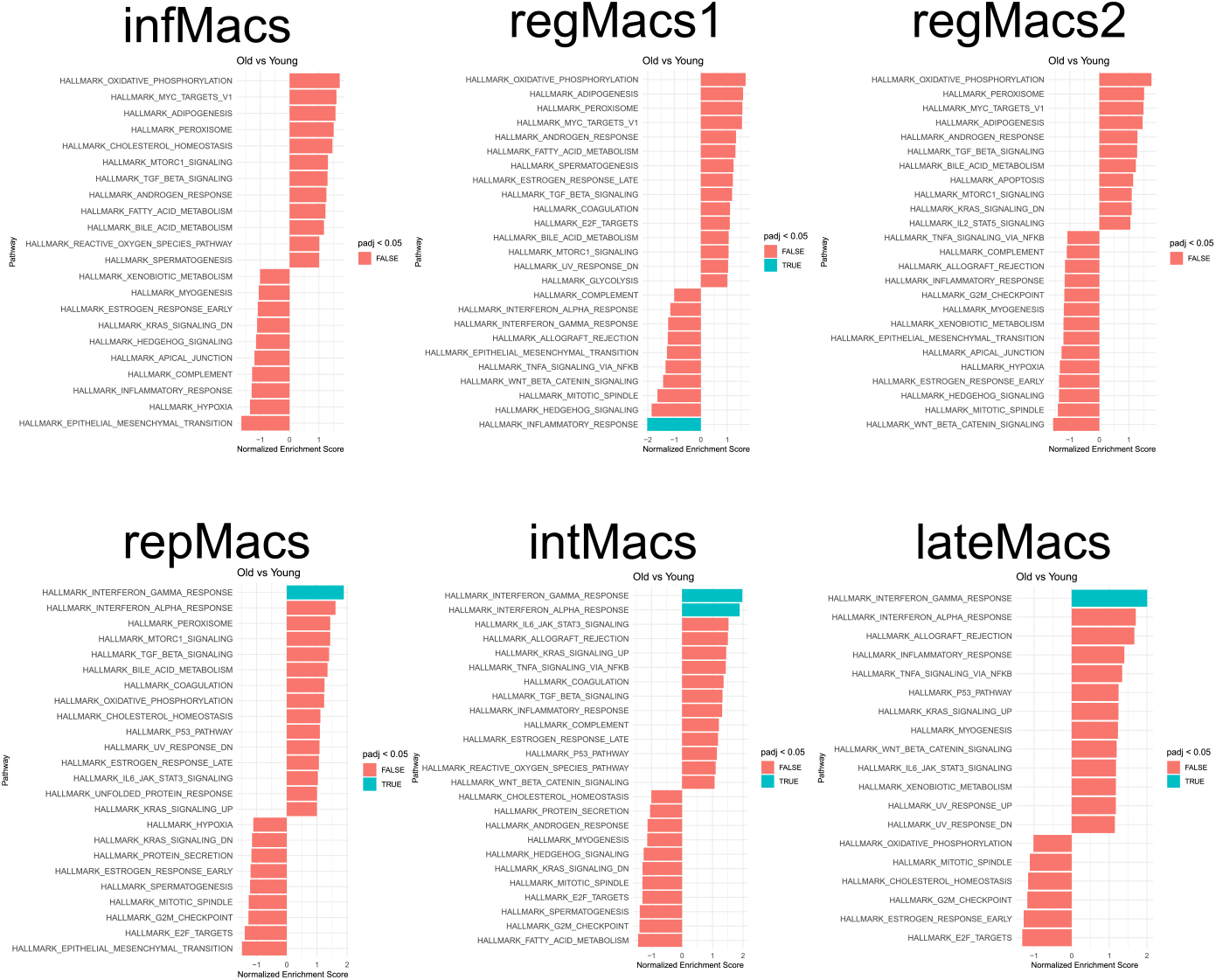
DisTwiner feature selection promotes distinct pathway enrichment profiles between cell clusters. Hallmark pathways GSEA scores plot for each cell cluster in the scRNA-Seq dataset using log-fold change from the DisTwiner-selected features comparing cells from aged and young samples as feature ranks.

## 4 Discussion

In this study we proposed an extension of the twiner classification model [20] that accounts for an arbitrary number of labels distributed across binary classification data, such as multiple disease variants in tumor data or different cell types in scRNA-Seq data. In addition, this study also allowed us to benchmark twiner performance with single-cell data and optimize the workflow accordingly. We initially defined a dissimilarity metric for DisTwiner by generating simulated data of a binary classification problem with multiple labels distributed across the two classes. We benchmarked multiple central tendency metrics of the coefficient distributions for DisTwiner optimization. The higher dispersion of feature coefficients computed by the minimum of the distribution in comparison to the mean or median increases the separation between features during model fitting, improving feature selection resolution and translating into coefficient computation being driven by pairs of labels with the most extreme dissimilarity measurements in the distribution. This aligns with the motivation of using DisTwiner to identify alterations of features that are characteristic of subsets of data samples. Additionally, we can further improve separation by applying an exponential transformation to the dissimilarity metric, which we believe to occur due to the exponential space increasing the smaller differences between dissimilarity measurements of a given feature.

Upon comparison of classification performance in scRNA-seq data, we observed that DisTwiner performed comparatively better than Random Forest classifying the young and aged cells in the scRNA-Seq dataset, but did not show significant improvements over Elastic Net classification, with both closing in on around 94% classification accuracy. These results are to be expected as `twiner`, and by extension DisTwiner, are derivations of logistic regression classification, with the most significant difference between the two models being the regularization term.

Parallel to the satisfactory classification performance of DisTwiner, another strong aspect of the model that our results highlight is the capacity to select features with heterogeneous fold-change profiles from control to test condition across all different class labels of the data samples. In particular, DisTwiner, and not the standard Elastic Net-regularized classifier, identified Cxcl13 as an important signal that is upregulated in aging primarily by intMacs, generating potential evidence regarding both the mechanisms of age-related expansions of the lymphoid compartment and the role of intMacs during regeneration. Comparing the dissimilarity weights of features selected by DisTwiner with that of features selected by Elastic Net also shows that DisTwiner tends to select features with low weights while the weights in the features selected by Elastic Net exhibits a tail of weights on the higher end of the distribution. Our results show that DisTwiner behaves as intended in the context of real transcriptomic data, selecting features for higher dissimilarity in the expression profiles of different data labels while preserving classification robustness of similar machine learning models such as Elastic Net.

We then assessed the features selected by DisTwiner for their biological significance in downstream functional analysis compared to features selected by either Elastic Net or MAST. We first observed a large overlap between features selected by DisTwiner and Elastic Net, whereas MAST not only selected a considerably different set of features but also selected a higher number of unique features in comparison to the other models. The higher number of features selected by MAST translated into a wider variety of pathway enrichments as GSEA showed. While exhibiting good classification robustness, logistic regression-based models such as DisTwiner and Elastic Net appear to not select a sufficiently high number of features capturing the biology of the GSEA hallmark pathways. Nevertheless, the GSEA results of DisTwiner versus Elastic Net suggest that the different features selected by the former method slightly improve the downstream GSEA analysis by identifying statistical significance in the IFN-*γ* pathway, which could only be achieved with Elastic Net by using it as an additional filter step downstream of MAST feature selection, thus a possible explanations can be that correlation-based penalization of features promotes the selection of biologically meaningful features or improves the filtering of features with excess noise that obscure result interpretation compared to standard regularization terms used in Elastic Net.

We then considered integrating the logistic regression-based models as an additional step of the MAST feature selection pipeline. By training the models with pre-selected features from MAST, we are adjusting the model fitting to a subset of features that are theoretically already labelled as important for classification, thus mitigating noise from non-selected features. This double feature selection additionally improves robustness of the output features, as we are selecting for features that pass more than one exclusion criteria. We ran GSEA on features selected by MAST + DisTwiner and MAST + Elastic Net combinations, and observed a narrowed down list of enriched pathways alongside alterations to the list of enrichments under the same thresholds such as the case of Kras signaling enrichment. We additionally observe when we compare Elastic Net feature selection with and without MAST-based pre-selection of features the emergence of a statistically significant pathway with the MAST preselection that the model did not detect by itself, suggesting that Elastic Net feature selection may improve in scRNA-Seq data by providing the model a more robust set of initial features with lower likelihood of carrying biological noise for the model to learn. These results suggest that logistic regression-based ML models, and by extension other ML models, serving as a second feature selection criteria in standard scRNA-seq analysis can improve selectivity in the downstream results, however both DisTwiner and Elastic Net perform similarly, so we lack evidence that DisTwiner feature selection is enough to tailor the functional analysis to a characteristic signature of each sample label.

DisTwiner and Elastic Net-selected features yielded less significant enrichments in gene set enrichment analysis compared to MAST-selected features without clear benefits in cluster-specificity, which may highlight a possible limitation of the classification model for the type of data used in this study. Both logistic regression-based classifiers selected a small set of features in comparison to the MAST differential expression test, thus this difference may originate in the base model selected and we do not exclude the possibility that DisTwiner may exhibit an adequate performance in data types other than scRNA-seq. Wani et al (2025) [25] performed a comparative study of multiple classification models in the context of scRNA-seq data, revealing differential performance between different model types, and that some models can for instance be more susceptible to the high dimensionality of the data or improper normalization/scaling, hindering their performance, which may explain the selection of different features by logistic regression models upon pre-selecting the features with MAST. Nevertheless, sophisticated ML models can be applied in the context of scRNA-seq data analysis, as Torang et al (2019) [26] demonstrated by constructing an Elastic Net-based model capable of discriminating multiple immune cell profiles in scRNA-seq data, but the behavior of this type of data begs careful consideration over the most appropriate models and data processing workflows in order to achieve successful results.

## 5 Conclusions

In this work we introduced DisTwiner, a generalization of the twiner correlation-based regularizer to an arbitrary number *K* of latent groups within a binary classification setting, and benchmarked it on scRNA-seq data of macrophage populations infiltrating young and aged regenerating murine skeletal muscle. By aggregating pairwise angular dissimilarities of feature correlation profiles across all group pairs, DisTwiner extends `twiner` beyond the two-group restriction while preserving its logistic-regression backbone. Benchmarking on synthetic and real data showed that the minimum-based aggregation combined with an exponential transformation yields the best trade-off between classification accuracy and separation of feature penalization weights, and that DisTwiner attains classification performance on par with Elastic Net and superior to Random Forest in this scRNA-seq setting. Feature-level inspection confirmed that DisTwiner preferentially selects genes with heterogeneous, cell-type-specific fold-change patterns that Elastic Net tends to discard, and functional analysis further indicated that regularized logistic-regression classifiers can act as a useful second-layer filter on top of differential expression tests such as MAST, refining the list of pathways carried forward to downstream biological interpretation. Although in the datasets analyzed here DisTwiner did not outperform Elastic Net in producing cell-type-specific functional signatures, its ability to natively accommodate multiple sample labels makes it a promising building block for future feature selection pipelines in single-cell and, more broadly, in high-dimensional multi-condition omics studies. Natural extensions include multi-class classification, adaptive weighting of group pairs based on sample size or biological prior, and integration with graph-based regularizers to jointly exploit correlation structure and known gene networks.

## 7 Data availability

The notebooks used for this study can be found at https://github.com/sysbiomed/DisTwiner. The scRNA-seq data used in this study is available at the NCBI Gene Expression Omnibus database under the accession number GEO: GSE279741

## Author Contributions Statement

T.M.C: Conceptualization, Methodology, Software, Validation, Formal analysis, Investigation, Data curation, Writing – original draft, Writing – review & editing, Visualization. M.B.L: Methodology, Writing – review & editing. J.N.: Resources, Writing – review & editing, Supervision, Project administration, Funding acquisition. P.S.-V.: Resources, Writing – review & editing, Supervision, Project administration, Funding acquisition. S.V.: Conceptualization, Methodology, Formal analysis, Writing– review & editing, Supervision, Project administration.

